# Kinase Inhibitor Pulldown Assay (KiP) for Clinical Proteomics

**DOI:** 10.1101/2022.10.13.511593

**Authors:** Alexander B Saltzman, Doug W Chan, Matthew V Holt, Junkai Wang, Eric J Jaehnig, Meenakshi Anurag, Purba Singh, Anna Malovannaya, Beom-Jun Kim, Matthew J Ellis

## Abstract

Protein kinases are frequently dysregulated and/or mutated in cancer and represent essential targets for therapy. Accurate quantification is essential, but current approaches are inadequate. For breast cancer treatment for example, the identification and quantification of the protein kinase ERBB2 is critical for therapeutic decisions. While immunohistochemistry (IHC) is the current clinical diagnostic approach, it is only semiquantitative. Mass spectrometry-based proteomics offers more quantitative assays that, unlike IHC, can be used to accurately evaluate hundreds of kinases simultaneously. The enrichment of less abundant kinase targets for quantification, along with depletion of interfering proteins, improves sensitivity and thus promotes more effective downstream analyses. Multiple kinase inhibitors were therefore deployed as a capture matrix for kinase inhibitor pulldown (KiP) assays designed to profile the human protein kinome as broadly as possible. Optimized assays were initially evaluated in 16 patient derived xenograft models (PDX) where KiP identified multiple differentially expressed and biologically relevant kinases. From these analyses, an optimized single-shot parallel reaction monitoring (PRM) method was developed to improve quantitative fidelity. The PRM KiP approach was then reapplied to low quantities of proteins typical of protein yields from core needle biopsies of human cancers. The initial prototype targeting 100 kinases recapitulated intrinsic subtyping of PDX models obtained from comprehensive proteomic and transcriptomic profiling. Luminal and HER2 enriched OCT-frozen patient biopsies subsequently analyzed through KiP-PRM also clustered by subtype. Finally, stable isotope labeled peptide standards were developed to define a prototype clinical method.

## INTRODUCTION

Protein and lipid kinases are enzymatic proteins that initiate and propagate signaling cascades to drive a wide range of cancer-relevant biological functions [1]. Many cancers are driven by aberrant kinase activity, and direct therapeutic inhibition of oncogenic kinases has proven to be effective for patients where individual driving kinases can be diagnosed – often through the presence of a genomic aberration [2, 3, 4]. The full scope of both regulation and downstream consequences of dysregulated kinase abundance and activity is not fully understood. Extensive crosstalk and functional redundancies of kinase-dependent dynamic signaling processes confounds therapeutic efficacy [5]. Consequently, while protein and lipid kinases inhibitors can successfully promote response and progression-free survival, achieving overall survival improvements has proved more challenging. The development of more effective kinome-based strategies therefore requires the accurate quantification of the kinome more broadly in human biopsy samples. Preclinical models can never cover the vast heterogeneity that exists in kinase function across cancers or link kinase expression to clinical outcomes.

Mass spectrometry-based comprehensive proteomics typically requires enrichment and/or prefractionation to effectively characterize low abundance kinases [6]. Deep-scale approaches achieve this goal through multiplexing with isobaric tags and pre-fractionation [7]. However, these approaches are cumbersome and impractical in a clinical setting where rapid data return is critical. Many kinases are not present in high abundance and are therefore more difficult to measure accurately. Furthermore, diagnostic clinical specimens present challenges in terms of the protein yields required for comprehensive kinome coverage. The development of methodologies to enrich kinases present in clinical samples is therefore also a critical endeavor.

Current enrichment strategies involve either antibodies or a kinobead approach [8, 9]. Immobilized kinase inhibitors have already been used to quantify low abundance kinases [10]. These approaches can utilize inhibitors with high specificity or with a broad affinity [11, 12]. As all type 1 kinase inhibitors are designed to bind the conserved ATP binding pocket domain of their targets, they can be used as an inhibitor-based affinity matrix for kinase enrichment in clinical samples. Extensive characterization of drug-kinase interactions has been performed with kinobeads [13], and specific kinobeads have been used to maximize kinome coverage [14]. However, these approaches typically use milligram quantities (∼5 mg) and their potential for tumor subtyping, characterization and risk-stratification has not been extensively evaluated [15].

In this study we developed a kinase inhibitor pulldown assay (KiP) with clinically relevant inhibitors that is optimized for microscaled quantities reflective of the yields of obtained from biopsy samples. We establish the coverage and quantitative fidelity of the assay for kinases in a single-shot discovery approach. From these data we optimize a one hundred kinase targeted panel and determined the effectiveness of KiP in subtyping breast cancer patient derived xerograph models and also two breast cancer patient sample cohorts.

## MATERIALS AND METHODS

### Kinase inhibitors

Palbociclib, Crizotinib, GSK690693 and AZD4547 were purchased from Selleckchem. Purvalanol B and CZC-8004 were purchased from Med Chem 101. Modified Afatininb, FRAX597, Abemaciclib, and Axitinib (containing an amino side chain for coupling) were custom synthesized by Med Chem 101.

### Kinobeads preparation

Kinase inhibitors Palbociclib, Crizotinib, GSK690693, AZD4547, CZC-8004, Afatinib, FRAX597, Abemaciclib and Axitinib were conjugated to ECH Sepharose 4B (GE Healthcare) via carbodiimide coupling chemistry as previously described [16]. For conjugation of the nine drugs with the reactive amine group, ECH Sepharose 4B (GE Healthcare) were used up to 2017, when this reagent was discontinued by the manufacturer. We synthesized our own ECH Sepharase 4B by conjugating 6-Aminohexanoic acid (Sigma) to cyanogen bromide (CNBr)-activated Sepharose 4B (GE Healthcare) according to manufacturer’s recommendation. Briefly, excess 6-Aminohexanoic acid was coupled to swollen CNBr-activated Sepharose 4B in 0.1M NaHCO3, pH 8.3 and 500mM NaCl at 4°C overnight with rotation. Unreacted CNBr groups were then inactivated by incubating the beads with 0.1M Tris-HCl pH 8.0 for 2 hours. The beads were then washed with five cycles of alternating low pH buffer (0.1M sodium acetate, pH 4.0 with 500mM NaCl) and high pH buffer (0.1M Tris-HCl, pH 8.0 with 500mM NaCl). Conjugation of the drugs to the “homemade” ECH Sepharose 4B were performed according to protocol described by Duncan et al [16]. Briefly, the beads were conditioned by multiple washes with 50% dimethyformamide/ethanol (DMF/EtOH). Each drug was dissolved in 50% DMF/EtOH and added to the conditioned beads in the presence of 0.1M 1-ethyl-3-(3-dimethylaminopropyl)carbodiimide (EDC) and allow to react overnight at 4°C with rotation. After coupling, unreacted groups were inactivated with 0.1M EDC, 1M ethanolamine in 50% DMF/EtOH for 1 hour at room temperature. Subsequently, beads were washed with 50% DMF/EtOH and alternating washes of 0.1M Tris-HCl, pH 8.3 with 500mM NaCl and 0.1M acetate, pH 4.0 with 500mM NaCl. Kinobeads were stored at in 20% ethanol at 4oC in the dark.

### Kinase enrichment by kinobeads precipitation (KiP)

For each KiP pulldown, 20 ug-200 ug of lysates were mixed with 10 uL of kinobeads that have been previously equilibrated in lysis buffer for 1 hour at 4°C with rotation. Kinobeads and its bound proteins were pulled down by centrifugation at 600x g for 30 seconds, the supernatant containing unbound proteins were aspirated. The beads were briefly washed with then successively washed two-times with 400uL buffer containing 50mM HEPES (pH 7.5), 600mM NaCl, 1mM EDTA, 1mM EGTA with 0.5% Triton X-100 and twice the same buffer without Triton X-100 followed by two washes with MS-grade water. After the final centrifugation, all the excess liquid was aspirated off and resuspended in 30 uL of 100 mM NH_4_HCO_3_ and heated at 65°C for 10 min. 2.5 ug of trypsin was then directly added to the beads and bicarbonate mixture and digested overnight at 37°C. To remove the remaining detergent prior to MS analysis, the digested peptide mixture was processed using the HiPPR Detergent Romoval Kit (Thermo) according to manufacturer’s directions and dried by speed-vac prior to MS analysis.

### Cell lines

Human melanoma cell lines SK-MEL-5 and MALME-3M were provided by Dr. Elizabeth Grimm (MD Anderson Cancer Center) and human leukemia cell line HL-60 and human prostate cancer cell line PC-3 were provided by Dr. Margaret Goodell and Dr. Jianming Xu (Baylor College of Medicine), respectively. Breast cancer cell lines T-47D and BT-474 were obtained from the American Type Culture Collection (Rockville, MD) and WHIM12 cell line was extracted from WHIM12 PDX tumor (Cell Rep, 2013, 24055055). SK-MEL-5 and MALME-3M cell lines were maintained in DMEM medium supplemented with 10% fetal bovine serum (FBS, Sigma-Aldrich, F2442) and HL-60, PC-3, and T-47D cells were cultured in RPMI-1640 supplemented with 10% FBS. BT-474 cells were cultured in DMEM/F12 medium supplemented with 5% FBS and 5 μg/ml insulin. WHIM12 cells were cultured in Ham’s F-12 medium containing 5% FBS with antibiotic and supplements (50 ng/mL sodium selenite, 50 μg/mL 3,3’,5-triiodo-L-thyronine, 5 μg/mL transferrin, 5 mM ethanolamine, 1 μg/mL hydrocortisone, 5 μg/mL insulin, 10 ng/mL Epidermal growth factor, and 2 mM L-glutamine).

### PDX tumor and cell lysates preparation

Frozen PDX tumors were cryopulverized with a Covaris CP02 Pulverizer and resuspended in lysis buffer containing 50mM HEPES (pH 7.5), 0.5% Triton X-100, 150mM NaCl, 1mM EDTA, 1mM EGTA, 1X protease inhibitor cocktail (Roche), 10mM NaF, 2.5mM Na_3_VO_4_ and 1% each of phosphatase inhibitor cocktails 2 and 3 (Sigma) and set on ice for 10 minutes prior to sonication. Cell lysates were sonicated in a Covaris S220 sonicator for 2 minutes at 4°C with settings at 100 peak power, 10 duty factor and 500 cycles/burst. Cell lysates were clarified by centrifugation at 100,000x g for 30 minutes at 4°C with a Beckman Optima Ultracentrifuge. Protein concentration were determined by Bradford assay (Bio-Rad).

For cell extracts, the cell lines were grown to approximately 80% confluency and harvested by scraping in cold PBS. The cell pellets were resuspended in lysis buffer and processed in the same way as the PDX tumor samples.

### Mass Spectrometry

#### Parallel Reaction Monitoring (PRM)

Digested peptides were analyzed by Orbitrap Fusion Lumos mass spectrometer coupled with EASY-nLC™ 1200 system (Thermo Fisher Scientific) for PRM. One fourth of peptides from KiP was loaded to a trap column (150 μm × 2 cm, particle size 1.9 μm) with a max pressure of 280 bar using Solvent A (0.1% formic acid in water), then separated on a silica microcolumn (150 μm ×5 cm, particle size, 1.9 μm) with a gradient of 5–28% mobile phase B (90% acetonitrile and 0.1% formic acid) at a flow rate of 750 nl/min for 75 min. Both data-dependent acquisition (DDA) and PRM mode were used in parallel. For DDA scan, a precursor scan was performed in the Orbitrap by scanning m/z 300–1200 with a resolution of 120,000 at 200 m/z. The most 20 intense ions were isolated by Quadrupole with a 2 m/z window and fragmented by higher energy collisional dissociation (HCD) with normalized collision energy of 32% and detected by ion trap with rapid scan rate. Automatic gain control targets were 5×10^5^ ions with a maximum injection time of 50 ms for precursor scans and 10^4^ with a maximum injection time of 50 ms for MS2 scans. Dynamic exclusion time was 20 seconds (±7 ppm). For PRM scan, pre-selected peptides were isolated by quadrupole followed by higher energy collisional dissociation (HCD) with normalized collision energy of 30% and product ions (MS2) were scanned by Orbitrap with a resolution of 30,000. Scan windows were set to 4 min for each peptide. For relative quantification, the raw spectrum file was searched with Mascot, and resulting mgf output was imported to Skyline with raw spectrum. Six strongest product ions were used to calculate peptide area. For accurate quantification, all AUC ranges were manually adjusted, and non-specific product ion was excluded. The sum of the area of product ions for each peptide was used to quantify each protein. Protein levels were median normalized, and log transformed for further analysis.

#### Peptide Synthesis

Peptides were purchased from Thermo Scientific Custom Peptide synthesis service and Vivitide. The peptides were purified to >95% purity by HPLC. Peptide mass was confirmed by mass spectral analysis and original concentration/net peptide content were determined by amino acid analysis. The peptides were dissolved in 20% acetonitrile/water and stored at −80C. A heavy peptide mixture stock was prepared by mixing an equimolar amount of each peptide with the concentration at 1 uM or 100 nM in 20% acetonitrile and 0.1% formic acid and it was diluted 10 times to make the final peptide mixture. Final peptide mixture was aliquoted to avoid multiple freeze/thaw and stored at −80C until use.

#### SureQuant

All SureQuant analysis was performed with Orbitrap Exploris 480 mass spectrometer (Thermo Scientific) coupled with Evosep One liquid chromatography system. Peptides were separated with 8 cm C18 column (EV-1109) using 30 samples per day method from Evesep One (44min gradient).

#### Survey MS analysis

Mixture of heavy peptides were separated with 8 cm C18 column (EV1109) using 30 samples per day method and the Exploris was operated in data dependent acquisition (DDA) mode with an inclusion list. Full scan spectra (300-1500 m/z, 120000 resolution) were detected by the orbitrap analyzer with automatic gain control targets of 300% and maximum injection time of 50 ms. For every full scan, up to 70 ions were subsequently isolated if the m/z was within +/-10 ppm of targets on the inclusion list and reached an intensity threshold of 1e^5^. Ions were collected with a maximum injection time of 10 ms, normalized AGC target 1000% and fragmented by HDC with collision energy 27% and detected with 150-1700 m/z and resolution 7500.

#### SureQuant analysis

Custom SureQuant acquisition template was built according to Thermo’s guidance and previously described method [17]. Full scan spectra were collected with 300-1500 m/z, AGC target 300%, maximum IT 50 ms, resolution 120000. Peptide matching the m/z within +/-3 ppm on the inclusion list were isolated (isolation window 1 m/z), fragmented (HCD collision energy 27%) and detected (150-1700 m/z, resolution 7500, AGC target 1000%, maximum IT 10 ms). A product ion trigger filter next performs pseudo-spectral matching, only triggering a MS2 event of the endogenous, target peptide at the defined mass offset if n>= 4 product ions are detected from the defined list. When triggered, light peptide MS2 scan was performed as follows: resolution 60000, 150-1700 m/z, AGC target 1000%, maximum IT 116 ms.

## RESULTS

### A combination of single-drug kinase inhibitors efficiently enrich the kinome

Nine different inhibitors (9KiP) were selected to maximize the coverage of kinases relevant for breast cancer: Palbociclib (CDK4/6 inhibitor), Crizotinib (c-MET and AXL inhibitor), CZC-8004 (non-specific tyrosine kinase inhibitor), Axitinib (VEGFR and PDGFR inhibitor), GSK690693 (AKT inhibitor), AZD4547 (FGFR and VEGF inhibitor), Afatinib (EGFR and ERBB2 inhibitor), Abemaciclib (CDK4/6 inhibitor), and FRAX597 (PAK inhibitor) (Figure 1A). Kinobeads were synthesized by coupling kinase inhibitors to ECH-sepharose beads via an amide bond (-C-N-). For this coupling reaction, the kinase inhibitor must contain a primary or secondary reactive amine (-NH_2_ and -NH, respectively) that can be covalently linked to the carboxyl group of the ECH-sepharose beads using EDC (1-ethyl-3-(3-dimethylaminopropyl)carbodiimide hydrochloride) as the reactive intermediate. As Afatinib, Abemaciclib, FRAX597 and Axitinib do not have the necessary amine reactive group, we modified these drugs to add a reactive group by adding a short C3 linker. In the case of Afatinib, an irreversible inhibitor of EGFR, we made an additional modification where the ethylene bond that reacts with Cys797 of EGFR was changed to an ethane bond. Removing the reactive ethylene bond on Afatinib derivative increases the ability of this drug to capture EGFR family members, including HER2, and other RTKs.

**Figure 1.**
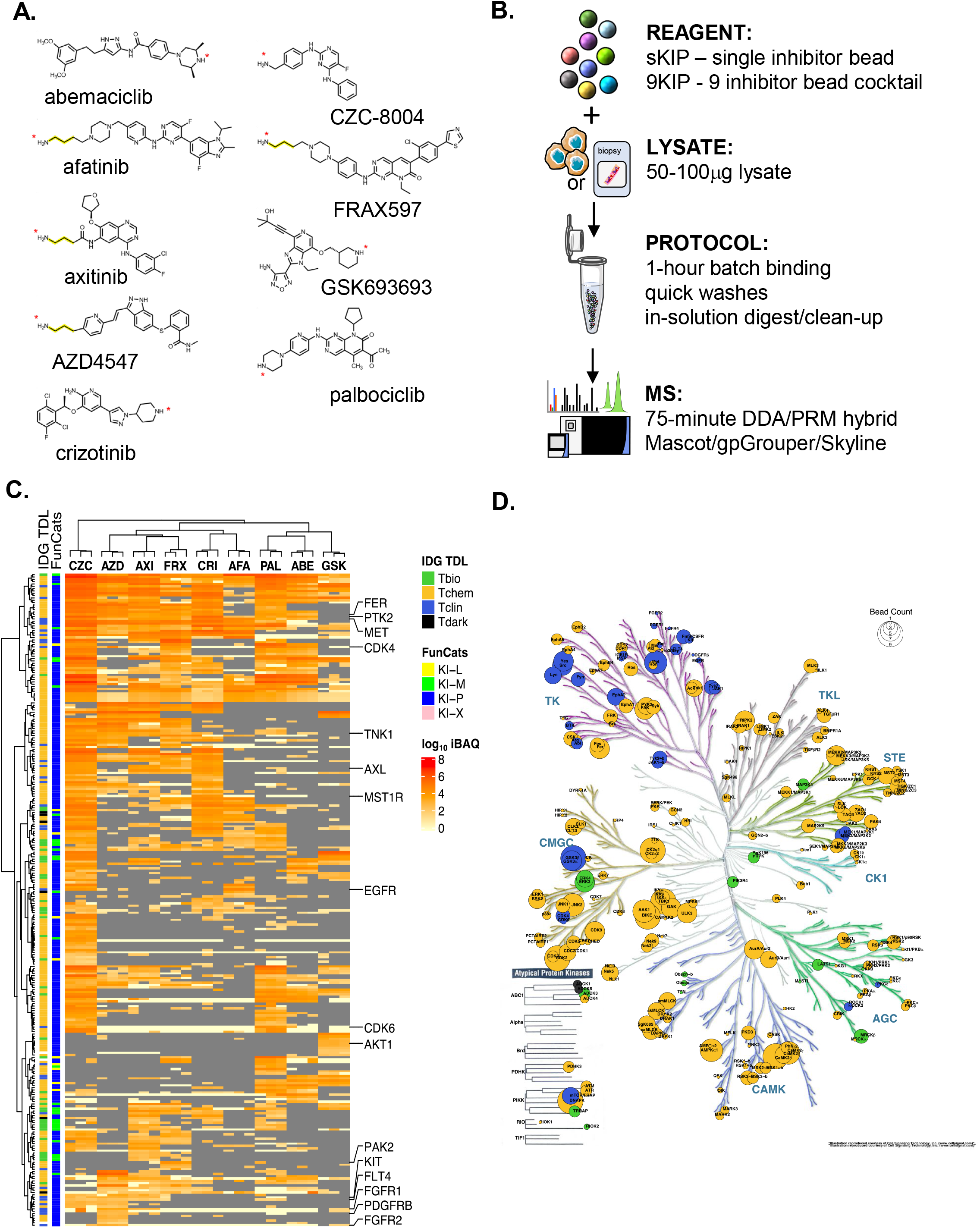
The kinase inhibitor pulldown assay. (A) Structure of 9 Kinase inhibitors used for KiP. For afatinib, axitinib, AZD4547, and FRAX597, a C3 Linker (yellow line) is added for conjugation. The amine group for conjugation is marked with an asterisk (*). (B) Workflow for the kinase inhibitor pulldown assay. Native protein lysates are incubated with kinase inhibitor-conjugated beads for 1 hour, and non-specific bound proteins are washed with high salt containing buffers. Inhibitor-bound kinases are digested with trypsin overnight, and digested peptides are cleaned with a detergent-removal kit and analyzed by mass spectrometry using a hybrid DDA/PRM mode. (C) Clustering of protein kinases enriched by individual inhibitors. Single inhibitor bead pulldown was carried out in triplicates for the 6 reference cell line mixture (6REF). Hierarchical clustering analysis of kinases in these experiments clearly shows that each kinase inhibitor pulls down distinct pool of kinases. Kinase classification by illuminating the Druggable Genome (IDG) is marked with different colors. (D) The Kinome tree with identified kinases highlighted. Colors represents IDG classifications, and the size of the circle represents number of inhibitors that can pull down that kinase.

We aimed to establish a microscaled kinase enrichment protocol that can ultimately be applied to diagnostic samples with limited material such as frozen tumor biopsies. There are two general approaches to enriching the kinome on immobilized inhibitor beads: large-scale column-based MIB (Multiplexed Kinase Inhibitor Beads) protocols that require milligram input scales and unpacked “batch” bead pulldowns such as those used by Kuster’s laboratory in 500ug scale. Since biopsies offer 20-100ug of protein for native protein lysates and clinical applications demand fast turnaround, we have focused exclusively on evaluating low input protocols. Our microscale KiP protocol greatly reduces the sample requirements for protein lysates to sub 50ug levels and can be completed in 2 days (Figure 1B).

### Characterization of single-drug kinase inhibitor pulldown beads (sKIPs) for kinome enrichment

A distinct spectrum of kinases was identified with each kinobead (Figure 1 C). To evaluate the immobilized inhibitors in capturing intended target kinases and additional off-target polypharmacology, we tested each individual kinobead-inhibitor using cell lysates. Because no individual cell type expresses every kinase in the genome, we used published RNA sequence data to identify six cell lines (6Ref) that together express a comprehensive array of protein kinases. Using 100ug of 6Ref lysate, each kinobead pulldown was performed in technical triplicates. For the eight substrate-specific kinase inhibitors (excluding the pan tyrosine kinase inhibitor CZC-8004), each were found to capture one or more of their intended targets, with exception of ERBB2, likely due to low levels of expression. ERBB2 capture was confirmed by adding the ERBB2-overexpressing BT-474 cell line to form the 7Ref mix used in all later protocol optimization experiments. In addition to protein kinases, we also detected significant number of metabolite and lipid kinases, attributed to the conservation of the ATP-binding domain. A high degree of reproducibility by label-free quantification was observed within each kinobead, with Pearson R values exceeding 0.9, underscoring the contribution of multiplexing drugs to maximum kinome enrichment (Supplementary Figure 1 A).

The kinobeads collectively cover a large spectrum of the human kinome (Figure 1 D). In addition to tyrosine kinase (TK) and tyrosine kinase-like (TKL) families, which are direct inhibitor targets, we observed kinases across all other families including Casein Kinase 1 (CK1) family, the serine/threonine kinase STE family, CMGC, AGC, Calmodulin/Calcium regulated kinases (CAMK), and some members of the Atypical Protein Kinase family. All kinobeads enrich kinases across more than one family (Supplemental Figure 1 B). While many kinases are overlapping between multiple kinobeads, each kinobead has a unique subset of kinases it can enrich (Supplemental Figure 1C). Importantly, most of the targetable kinases with a clinically approved drug (T-clin) or with a pre-clinical tool compound (T-chem) by the NIH Illuminating the Druggable Genome (IDG) Consortium, are well represented. This highlights the potential of our kinome profiling approach to uncover unexpected therapeutic avenues where immediate application or repurposing can be indicated based on clinical sample profiling.

### Microscaled KiP efficiently enriches the kinome

To further investigate the feasibility of microscaling the assay to protein yields typical of core biopsy specimens, we performed kinome pulldowns across a range of protein inputs with the 9KiP reagent and the 7Ref lysate. For assessment of assay linearity in low-microscale samples, 10 uL 9KiP beads were used with increasing amounts of 7Ref lysate (0, 12.5, 25, 50, 100, and 200 μg) (Figure 2 A). More than 220 kinases were identified at the lowest level of 12.5 ug lysate, increasing to ∼300 kinases with 50 ug of input (Figure 2 B). Increasing input did not further increase the number of kinases observed, but total ion current and the total intensity of kinases detected increased linearly (Supplementary Figure 2 A). 50 ug of protein lysate is sufficient input for the MS-based detection of the expressed kinome after KiP enrichment. The total amount of bound kinases, evaluated here as total iBAQ value, scaled linearly across all levels of input, suggesting that 10 uL of our 9KiP cocktail is not saturated with kinome from up to 200 ug of lysate, and the KiP assay has a good linear range for kinome recovery for low microgram protein samples. The linear relationship to protein input holds true for most identified kinases, including ERBB2 (Supplemental Figure 2 A). Quantification of some highly abundant kinases, including CDK4 and PRKDC, begins to saturate at 50 ug lysate (Supplemental Figure 2 B-E). These data underscore the importance of characterizing individual kinase response curves should accurate quantification be needed in more stringent clinical setting.

**Figure 2.**
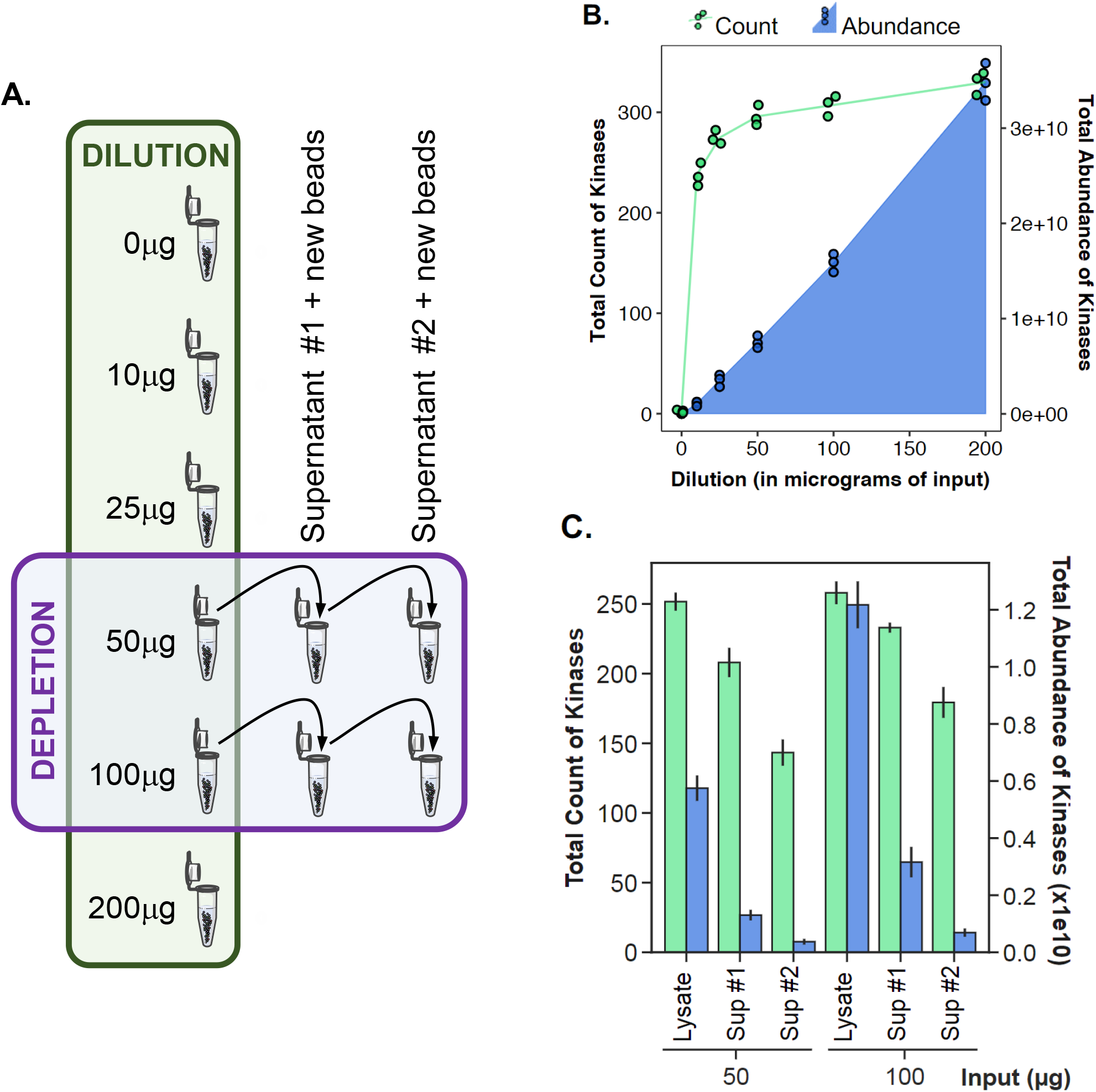
Linearity and binding efficiency of KiP. (A) Schematic for KiP with different input amounts and serial depletion. KiP was performed with increasing amounts of 7REF lysate. For the depletion experiments, KiP was repeated twice with the supernatant from 50ug and 100ug KiP experiments. (B) Number of kinases identified and kinase abundance from different input experiments. Kinase numbers are plotted in green and abundance is plotted in blue. (C) Number of kinases identified and kinase abundance from depletion experiments. Kinase numbers are plotted in green and abundance is plotted in blue.

To determine the binding percentage and capacity of KiP, pulldowns were repeated with supernatants containing unbound proteins from preceding KiP enrichments (Figure 2 A). 10uL of 9KiP beads were used with 50ug and 100ug amounts of 7Ref lysate, and their supernatants were subjected to two subsequent rounds of KiP enrichment with new beads. In each round, kinases decrease modestly by number of identifications but dramatically by level (Figure 2C). With 50ug of lysate, 84% of kinases are identified after the first depletion, and 59% of kinases after the second. The first depletion input recovers 28% of the original kinase abundance, and the second depletion less than 10%. These trends were similar for 100ug, however more kinases were observed overall as total levels were higher than those observed for 50ug. The total kinases observed, linear range and binding percentage are sufficient to reliably utilize the pulldown to quantify kinases. The data dependent acquisition schemes utilized for these experiments are stochastically less quantitative for lower abundance kinases. To increase quantitative metrics, we optimized a parallel reaction monitoring (PRM) assay with targets identified in our pulldown assays.

### PRM assay development and parameters

KiP-PRM yields sensitive, accurate and precise measurements of kinases. To improve the sensitivity, accuracy, and precision of the assay, we designed PRM assays with representative peptides across 54 druggable kinases and 46 kinases found to be differentially expressed in previous profiling experiments of breast cancer patient derived xerograph models for a total of 100 kinases. Initially, five to seven peptides were chosen empirically based on their performance in a collection of all experiments from our laboratories (>7000 experiments, Supplemental Figure 3A) and filtered by being unique-to-human, unique-to-gene and representative of all gene-specific isoforms, when possible. We tested linear responses of these peptides on dilutions of the same KiP lysate and identified 2-4 peptides by best response curve and peptide peak shapes (Supplemental Figure 3B).

Kinases are quantified by PRM with equal or greater precision than when quantified by DDA. KiP experiments for 7Ref lysate were performed across 4 different concentrations (12.5 μg to 100 μg) in technical duplicates, and samples were run on the mass spectrometer with a 75 min label-free hybrid DDA/PRM method including >200 PRM targeted peptides with 5-minute retention time windows. All proteins show very strong correlation to input levels from PRM quantification, regardless of their relative abundance (Figure 3A). In contrast, while DDA achieved good correlation for the most abundant kinases, measurements deteriorated for lower-level proteins. For example, both DDA and PRM show consistent quantification for ERBB2, the 16th most abundant kinase in intensity (Figure 3B). However, the less abundant CDK6 kinase was not consistent by DDA, whereas its levels by PRM followed input levels accurately. Poor correlation for less abundant kinases by DDA is partly due to lack of peptide identification; however Pearson correlation coefficients are better from PRM than from DDA even when samples without identifications are excluded from analysis (Figure 3B). This suggests that PRM quantification with verified peptides markedly improves quantification accuracy of enriched kinases, particularly those of lower abundance.

**Figure 3.**
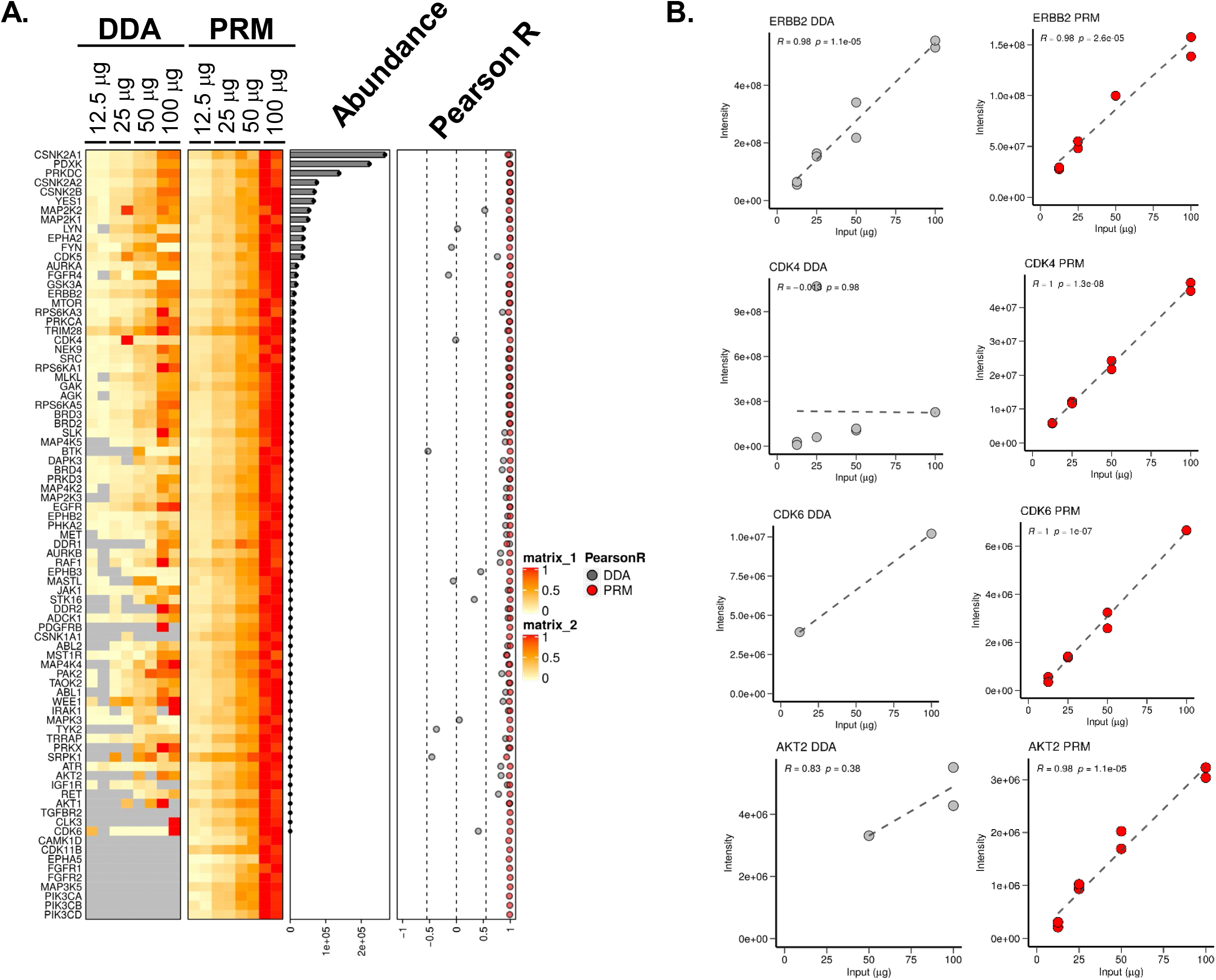
Quantification of kinases by DDA and PRM. (A) KiP experiments for 7REF cells were performed at 4 different concentrations (from 12.5 μg to 100 μg), and samples were run on the mass spectrometer using hybrid mode (DDA/PRM). Although both DDA and PRM produce good correlations for many kinases, PRM dramatically improves quantification for lower abundance kinases. (B) ERBB2, CDK4, CDK6, and AKT2 quantifications are plotted as representative examples.

### PDX subtyping with KiP-PRM

KiP-PRM classifies and subtypes breast cancer xenografts in concordance with comprehensive molecular profiling. We applied KiP to 16 breast cancer xenografts previously generated and characterized by deep transcriptome, proteome, and phosphoproteome sequencing [18]. We previously demonstated that these models are relatively stable across passages and thus are a representative set of breast cancer subtypes [19]. 50 μg of protein lysate from PDX tumors was used for KiP enrichment, and all experiments were performed in technical duplicates. One third of peptides post KiP enrichment were analyzed using a DDA/PRM hybrid method, and a total of 91 kinases were quantified by PRM. Hierarchical clustering using kinases measured by PRM separates tumors into groups corresponding primarily to basal and luminal PAM50 subtypes (Figure 4). The two HER2-expressing tumors (WHIMS 8 & 35) fall into the two different main clusters, suggesting substantial kinase differences between the models. Nevertheless, both tumors have the highest expression of ERBB2 across the cohort, as expected. Lastly, the claudin-low tumor WHIM12 clusters near, but distinct from the basal subgroup. This is consistent with the consensus that claudin-low tumors can be related to basal subtypes [20] and with a recent finding that most claudin-low tumors have a basal-like intrinsic subtype [21].

**Figure 4.**
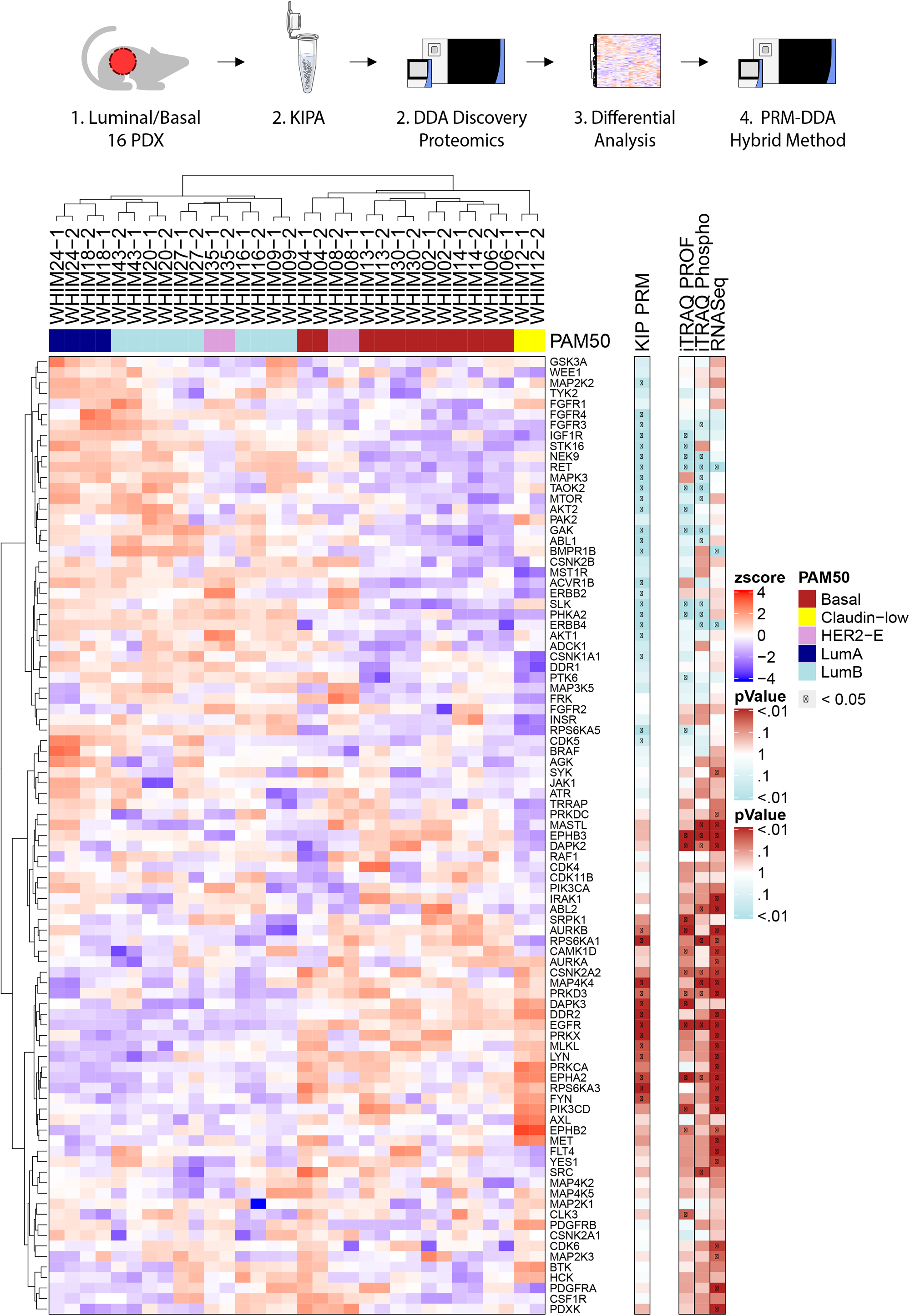
KiP classifies 16 WHIM PDX tumors according to their intrinsic subtype. Kinases from 16 breast cancer xenografts were enriched by KiP, and druggable kinases and subtype-specific kinases were quantified by PRM. 50 μg of protein from PDX tumors was used, and all experiments were performed in duplicates. Clustering analysis of kinases distinguishes basal subtype samples from luminal subtype samples, and claudin-low samples are separated from all other samples. A basal specific kinase, EGFR is enriched in basal PDXs, and luminal specific kinases, RET and IGF1R, are enriched in luminal PDXs. KIP is able to capture most of subtype-specific kinases identified in previous RNA-sequencing or proteome profiling studies (Huang, K.-l. et al. Proteogenomic integration reveals therapeutic targets in breast cancer xenografts. Nat. Commun. 8, 14864 doi: 10.1038/ncomms14864 (2017)).

To approximate the statistical power of the KiP-PRM assay, we performed two-sample t-tests between the basal and luminal subtypes (grouping luminal A and B together) for all proteins that were quantified by the assay as well as for analytes from the previously published iTRAQ protein, iTRAQ phosphoprotein, and RNASeq datasets [18]. The mean value in each technical replicate was used for calculation for the KiP-PRM data. From the KiP data, using the uncorrected p-value threshold of 0.05, 36 kinases are significantly differentially expressed between PDX models. 13 Kinases are upregulated in basal models, and 23 kinases are upregulated in luminal. As expected, EGFR is enriched in basal PDXs, and known luminal BRCA associated kinases RET and IGF1R are elevated in luminal PDXs.

KiP differentially quantifies more subtype specific kinases (36 kinases) than iTRAQ proteome profiling (24 kinases). This additional quantification fidelity likely arises from enrichment and increased precision due to PRM targeting: most significant kinases in the KiP-PRM data have similar directionality in the proteome and phosphoproteome dataset. Further, KiP-PRM measurements correlate well across the majority of proteins with a median Pearson correlation above 0.5 for both protein-based measurements - in contrast the correlation with RNASeq which exhibits a bimodal distribution and a lower median Pearson correlation of 0.23.

RNA-sequencing identifies a similar number of subtype-specific kinases (35) that poorly overlap with those found to be significant in proteomic profiling or KiP. The distribution of significant kinases from the RNA-seq data is uneven with 32 being basal specific kinases while only 3 are upregulated in luminal tumors. Interestingly, when examining all proteins identified through KiP-DDA, many RNA related proteins appear upregulated in basal BRCA. These results are consistent with previous comparisons between RNA and protein expression levels [22]. While there is substantial overlap with previous data with regard to subtype-specific proteins, KiP PRM illuminates additional kinases that differ between subtypes not necessarily exhibited from previous omics analyses. This reflects the enrichment and precise quantification afforded by KiP-PRM.

### Patient sample subtyping with KiP-PRM

Given the subtyping of PDX tumors achieved using KiP, we pursued the analysis of clinical samples obtained from previously studied cohorts: Luminal subtype from Preoperative Letrozole (POL) clinical trial [23], and ERBB2+ samples from the Discovery protocol 1 (DP1; NCT01850628) study. Approximately 50-100 ug of lysate obtained from OCT-frozen blocks was used for each enrichment, and one-third to one-half of the post-enrichment protein pool was used for analysis. The majority of samples analyzed have previous RNAseq results, from which stromal scores, immune scores and their combination: Cibersort, xCell and ESTIMATE scores were derived [24]. These metrics inform of the quality and purity of the tumor sample and the microenvironment of the tumor. In total, 16 luminal and 21 ERBB2+ (HER2) samples were analyzed [Figure 5]. K-means clustering of the samples yielded 3 subgroups: one Luminal and two HER2 groups. Resistant and sensitive samples tend to group together within subtypes, and ESTIMATE scores were highest in HER2 cluster 3.

**Figure 5.**
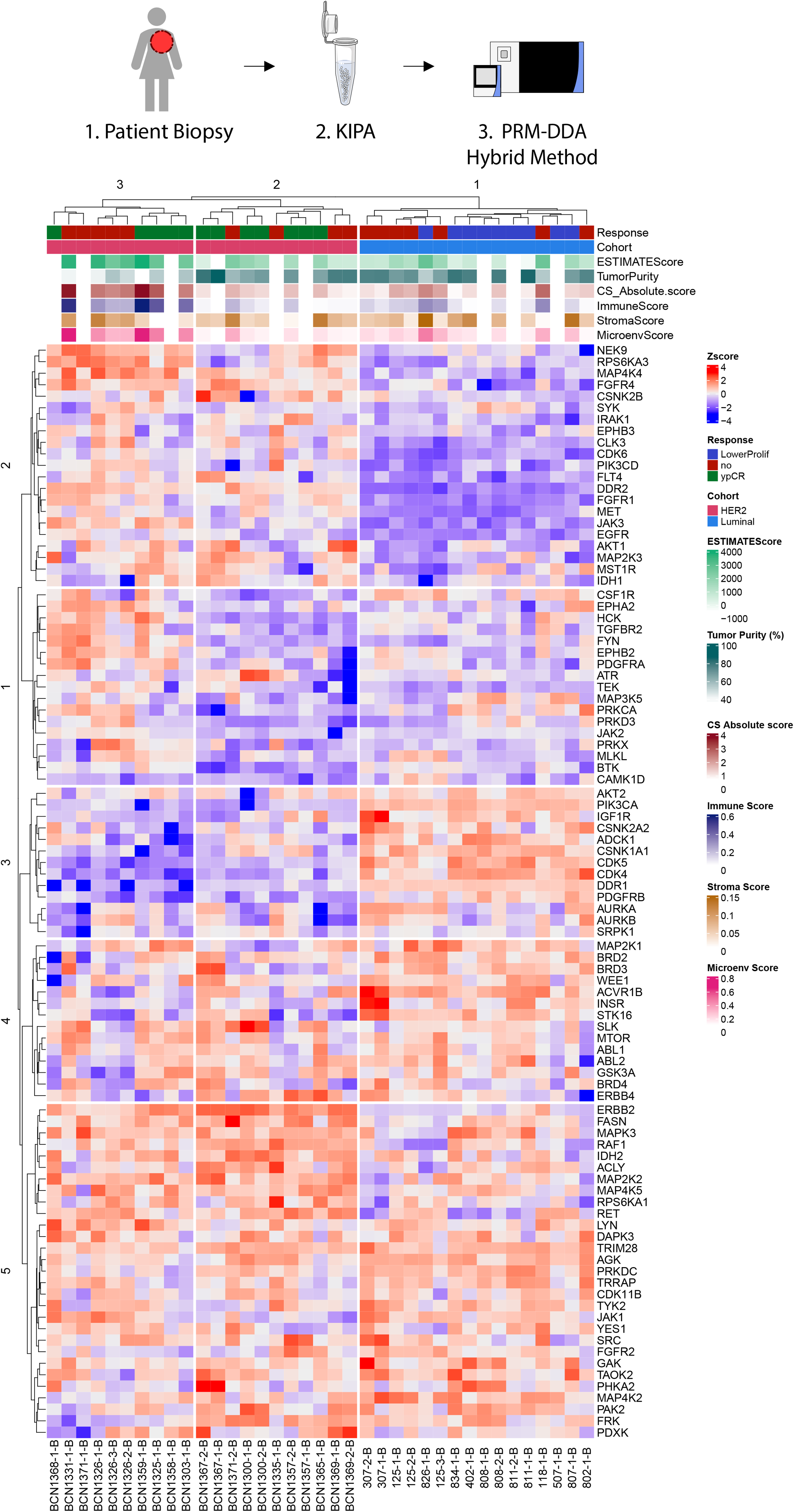
KiP clusters breast cancer patient samples by subtype by quantifying kinases. KiP-PRM clusters breast cancer patient samples by intrinsic subtype, identifies subgroups within subtypes, and partially clusters resistant patients within subtype. 37 patient samples from a HER2+ cohort and a luminal cohort were processed with the KIP assay and analyzed by PRM. HER2 is highly enriched in the HER2+ cohort, and luminal associated kinases such as CDK4 were elevated in the Luminal cohort.

HER2 is significantly elevated in the HER2 cohort (p-value = 8E-10 Welch’s t-test) with a 53.89-fold change as compared to the luminal cohort. DDR2, FGFR1, JAK3, MAP4K4 and CSNK2A1 are also elevated in the HER2 cohort, with DDR2 being the best delineator of the two subtypes with a p-value of 7E-18 and 188.64 fold change versus luminal. Conversely, CDK4, CDK5 and PIK3CA were significantly elevated in the Luminal cohort (p-value < 1E6 Welch’s t-test). These findings are consistent with previous ER+ and HER2+ breast cancer studies and our observations with PDX models. Overall KiP-PRM succeeded in clustering breast cancer patient samples into clinical subtypes using 100 ug of lysate or less.

### IS-PRM subtyping with KiP

Encouraged by the effective subtyping of patient samples with KiP, we decided to pursue a clinically applicable absolute quantification approach. To maximize clinical efficacy of our KiP approach, we developed an Internal Standard Triggered-Parallel Reaction Monitoring (IS-PRM) assay using 106 heavy peptides identified from our PRM studies. We removed the hybrid component of the PRM method and shortened the gradient to 44 minutes, resulting in throughput of 30 injections a day. Lysates used in the previous PDX-PRM analysis were combined with either 10fmol (98 peptides) or 100fmol (8 peptides) of stable isotope labeled peptides. All results were manually validated using Skyline. To establish the quantitative parameters of this approach, we ran a mixture of Luminal PDX samples in 18 replicates. We achieved an average coefficient of variance (CV) of 9.28% (2.2%-27.2%) for 80 peptides (Supplemental figure 3). IS-PRM KiP recapitulates the clustering observed in the KiP-PRM data. Luminal and basal PDX models cluster together [Figure 6]. Further, Luminal A, Luminal B and HER2 subtypes clustering is comparable with that observed in the PRM dataset.

**Figure 6.**
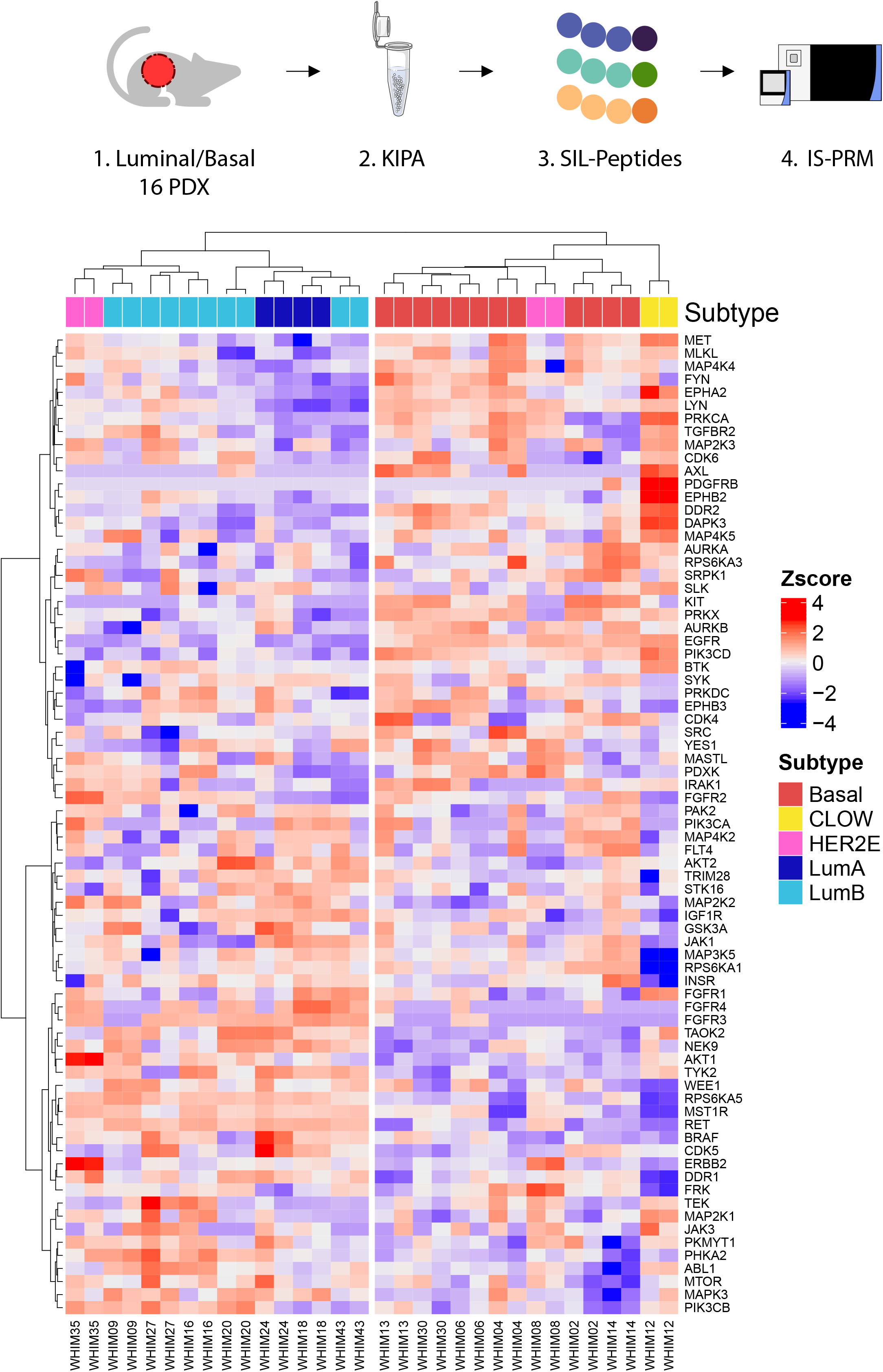
Prototype clinical implementation with IS-PRM for 16 PDX KiP samples. Heavy isotope labeled triggered acquisition of KiP enriched kinases in combination with an evosep and Exploris 480 as a prototype clinical assay. Nearly identical clustering observed in the PRM method was obtained.

## DISCUSSION

The kinase inhibitor pulldown assay is a robust, reproducible and clinically relevant approach tosuccessfully enrich and quantify the majority of the kinome.. The current inhibitor combination used in our pulldown yielded over 300 distinct kinases, and it is likely that additional kinases of interest can be identified through further assay development. Should a novel inhibitor target a kinase that is not binding to the current KiP assay, that novel inhibitor may be immobilized and added as an additional component to the matrix. During the development of this method, we attempted multiple different inhibitor combinations and found no detriment to identification capacity with additional inhibitors. Thus, KiP is modular and additional inhibitors may be added. This capability may even extend beyond kinases and into other low-abundance and biologically relevant targets.

To our knowledge, this study presents the first classification of breast cancer PDX and patient samples solely on kinase abundances. While the patient sets are from vastly different cohorts, and thus expected to have analytical differences, KiP identified kinases with an extreme dynamic range afforded by relatively clean, and abundant spectra due to enrichment. This is most obvious in the IS-PRM dataset where subtype defining kinases are orders of magnitude differentially expressed between samples. In contrast, isobaric labeling approaches failed to identify kinases as differentially expressed as consistently PRM did. With the experimental design of this study, it is impossible to know what the ‘base truth’ is for differences in kinase expression between samples, and thus KiP may have yielded false positives. However, given the well-established phenotypic and pathological differences between luminal and basal breast cancer, there is potential utility in any differentially identified target.

In this study Orbitrap instruments, DDA and PRM/IS-PRM were exclusively used with KiP. The benefits of KiP enrichment are mass-spectrometer universal, and other available platforms with triple quadrupole and time-of-flight detectors will also have empirically improved quantification of kinases. Additionally, data independent acquisition techniques will similarly benefit as spectra will inherently be less convoluted. Similar benefits are observed with antibody-based enrichment. A critical step towards non-antibody-based enrichment in a clinical setting is extensive validation.

A major challenge in the clinical implementation of KiP is the need for native lysate. Effective binding of kinases to inhibitors required kinases to not be denatured or cross-linked. Current methods of extracting proteins from FFPE blocks use heat, xylene and/or detergents. These methods are not compatible with KiP. Flash frozen and OCT-embedded samples are viable alternatives, yet their clinical implementation is not widespread. An efficient protocol for accrual and banking of these samples is necessary for further progress.

## Acknowledgements

Authors would like to acknowledge funding support CPTAC PTRC grant U01CA214125 (MJE/MA, SAC), National Cancer Institute’s Specialized Programs of Research Excellence (SPORE).

BCM Mass Spectrometry Proteomics Core is supported in part by the Dan L. Duncan Comprehensive Cancer Center Award (P30 CA125123), CPRIT Core Facility Awards (RP170005 and RP210227).

Authors would like to acknowledge the intellectual contributions from Shankha Satpathy, Ph.D. and Steven A. Carr, Ph.D.Ethical Compliance:

## FIGURE LEGENDS

**Fig S1.**
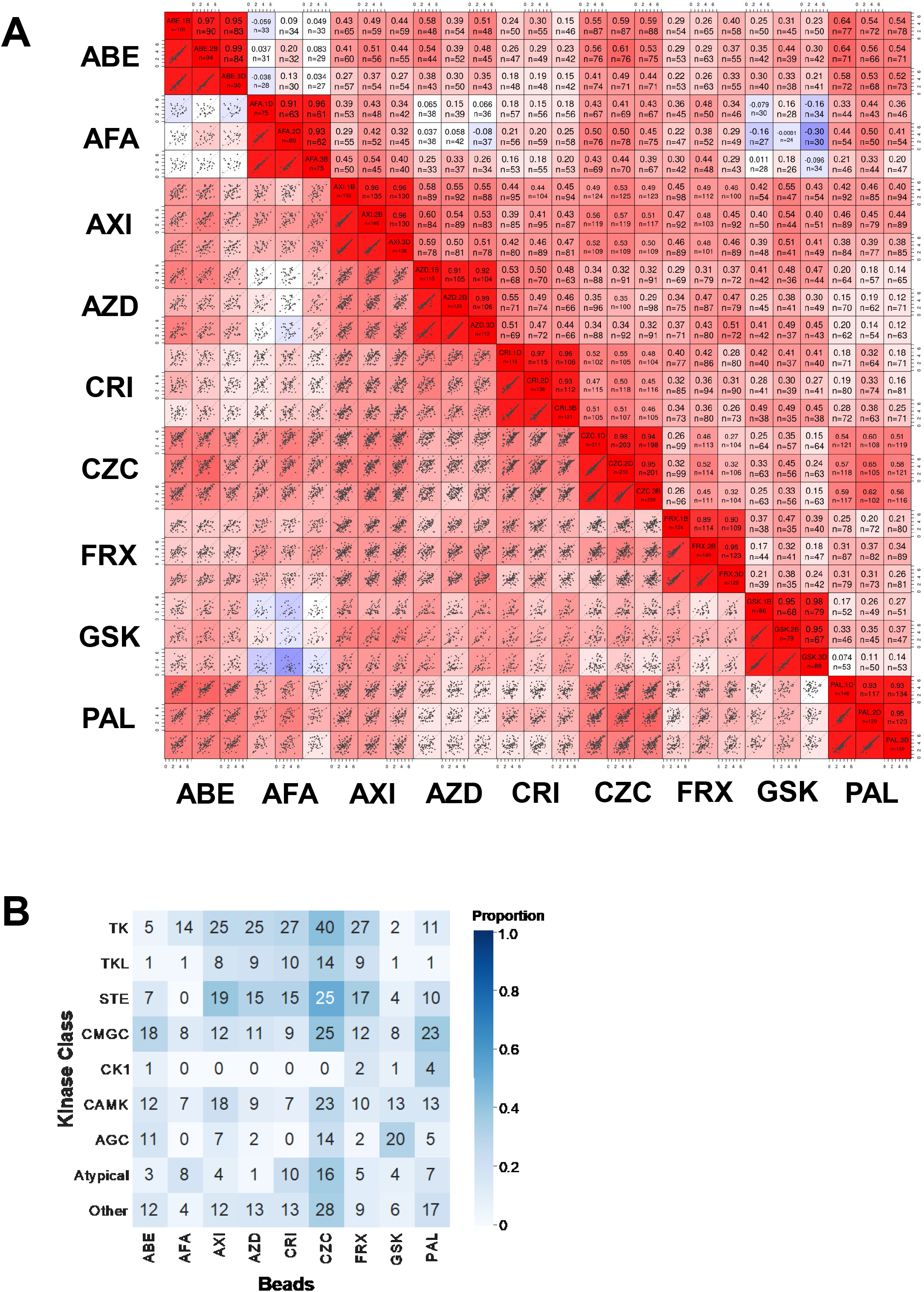
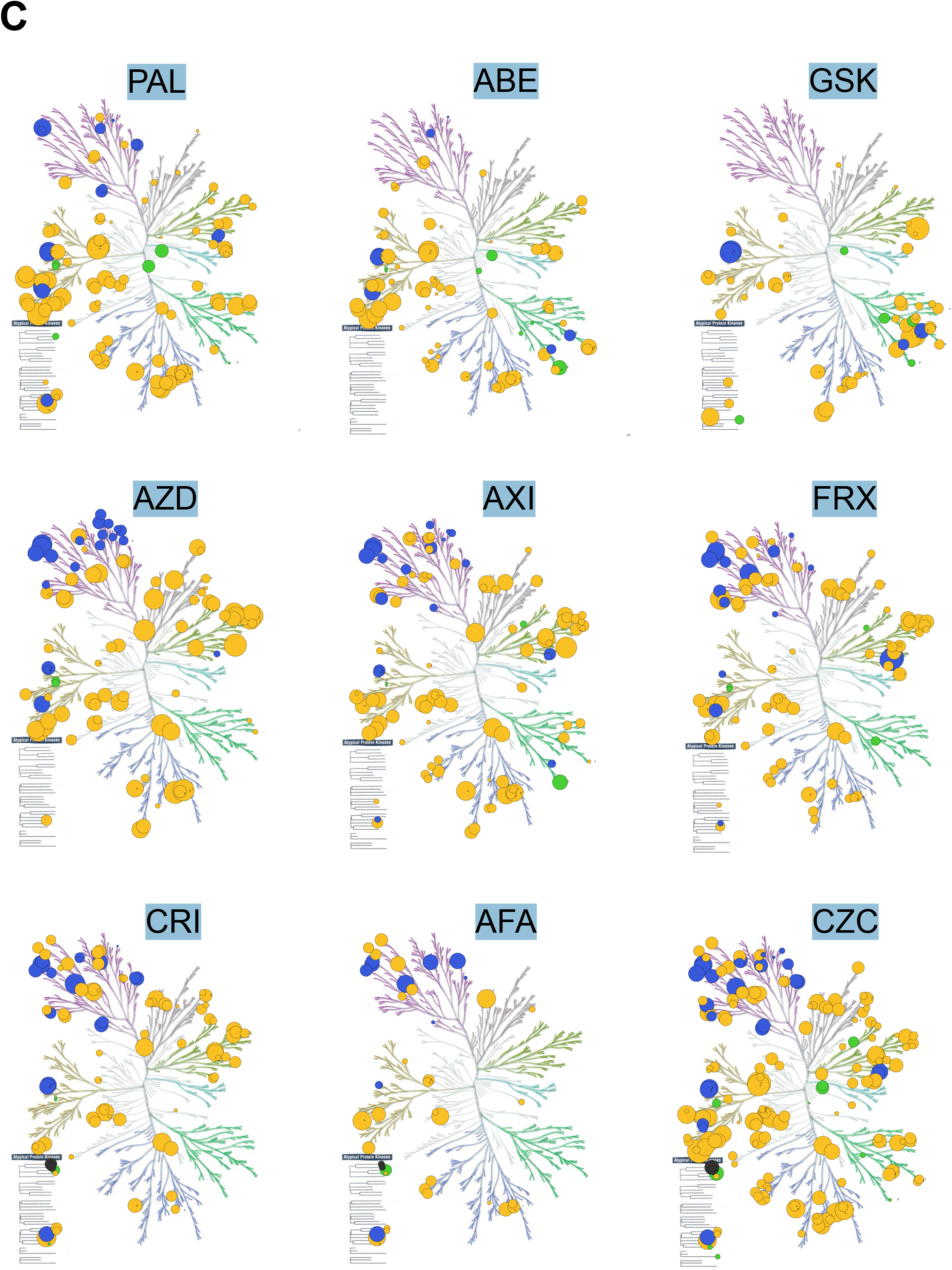
KiP with single inhibitor beads (sKiP) **(A)** Repeatability KiPs by 3 different technicians are reproducible. **(B)** Kinase family identified by sKiP **(C)** Kinome Tree by sKiP

**Fig S2.**
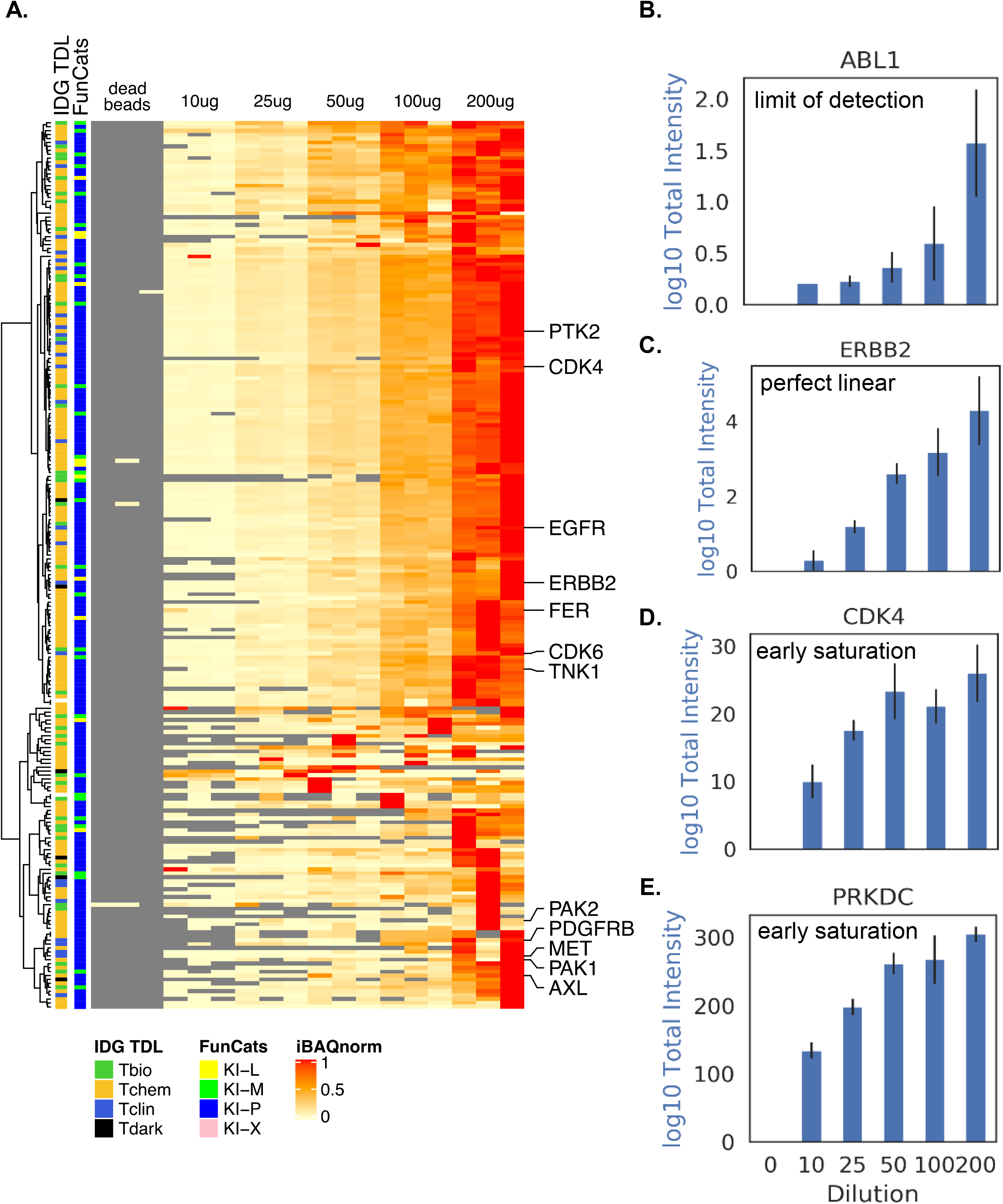
KiP with different input experiment. KiP was carried out with different amounts of lysate (Figure 2A) and quantified kinase levels are plotted (A). (B-E) ABL1, ERBB2, CDK4 and PRKDC quantification was plotted as representative examples.

**Fig S3.**
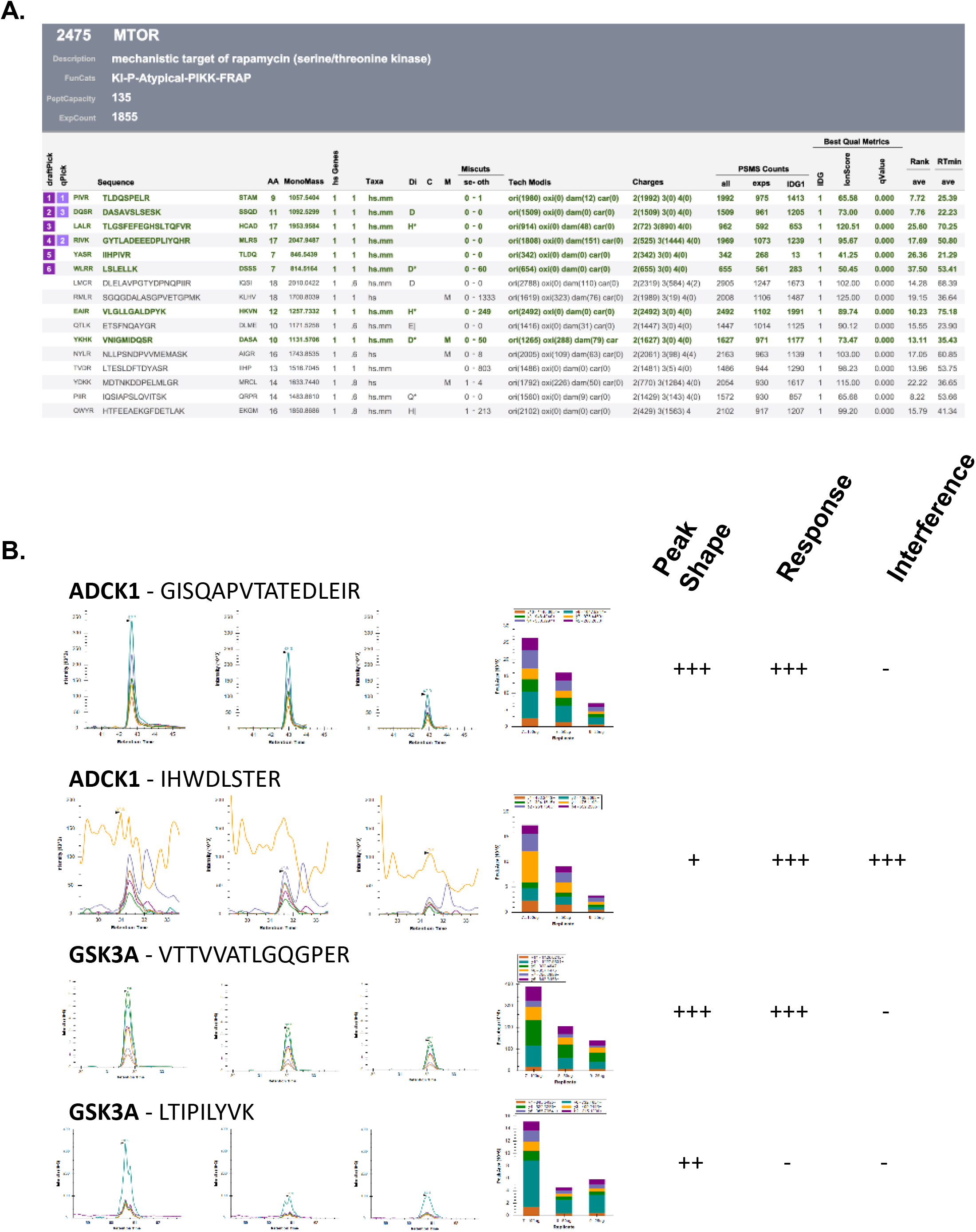
qPick. (A) Example of **qPick** from **iSPEC** database All the information of identified peptides for each kinase in iSPEC database are presented by qPick. It includes peptide sequence, mass, gene product number for the peptide, miscleavage, PSMs for each modification, PSMs for each charge, best ion score, average retention time, etc. Peptides were ranked by experimental PSM counts and top 3 to 6 peptides were selected for PRM runs. However, peptides are possibly excluded if they fall into following categories; More than 10% of PSMs has modification (2) A peptide has miscleavage in it (3) Many PSMs of that peptide are part of miscleaved peptides (4) Sequence is shared with other gene product (5) bad ion score (< 20). (B) Representative examples of PRM peptide selection KiP experiment was performed with different amount of inputs and samples ran on mass spectrometry with PRM method to choose best PRM peptides. We took following categories into consideration. (1) Peak shapes – peaks need to be symmetrical and narrow (2) response – sum of peak areas should be proportional to input level (3) interference – there should be no other non-specific peaks. We chose the peptides which meet these categories and generated the final list of PRM peptides for kinases. For example, peptide IHWDLSTER for ADCK1 has non-specific peaks around although peptide response looks good. On the other hand, peptide LTIPILYVK for GSK3A shows bad response whereas peak shape is good and there is no non-specific band. Therefore, these peptides had not been selected as final PRM peptides.

